# Short-term circulating tumor cell dynamics in mouse xenograft models and implications for liquid biopsy

**DOI:** 10.1101/814368

**Authors:** Amber L. Williams, Jessica E. Fitzgerald, Fernando Ivich, Eduardo D. Sontag, Mark Niedre

**Author notes:** Corresponding author: Mark Niedre, Northeastern University, 360 Huntington Avenue, Boston, MA, USA, **Email:**.

## Abstract

**Motivation:** Circulating tumor cells (CTCs) are widely studied using liquid biopsy methods that analyze single, fractionally-small peripheral blood (PB) samples. However, little is known about fluctuations in CTC numbers that occur over short timescales *in vivo*, and how these may affect accurate enumeration from blood samples.

**Methods:** We recently developed an instrument called ‘diffuse in vivo flow cytometry’ (DiFC) that allows continuous, non-invasive counting of rare, green fluorescent protein expressing CTCs in large deeply-seated blood vessels in mice. Here, we used DiFC to study short-term changes in CTC numbers in multiple myeloma and Lewis lung carcinoma xenograft models. We analyzed 35- to 50-minute data sets, with intervals corresponding to approximately 1, 5, 10 and 20% of the PB volume, as well as changes over 24-hour periods.

**Results:** For rare CTCs, the use of short DiFC intervals (corresponding to small PB samples) frequently resulted in no detections. For more abundant CTCs, CTC numbers frequently varied by an order of magnitude or more over the time-scales considered. This variability far exceeded that expected by Poisson statistics, and instead was consistent with rapidly changing mean numbers of CTCs in the PB.

**Conclusions:** Because of these natural temporal changes, accurately enumerating CTCs from fractionally small blood samples is inherently problematic. The problem is likely to be compounded for multicellular CTC clusters or specific CTC subtypes. However, we also show that enumeration can be improved by averaging multiple samples, analysis of larger volumes, or development of new methods for enumeration of CTCs directly *in vivo*.

## 1. Introduction

Circulating tumor cells (CTCs) are of great interest in cancer research because of their importance in hematogenous metastasis. CTCs shed from the primary tumor into the peripheral blood (PB), and a small fraction may form metastases. It is these metastases that are extremely difficult to control clinically and eventually result in the majority of cancer-related deaths (1, 2). Nearly all CTC clinical and pre-clinical research involves “liquid biopsy”, wherein CTCs are isolated from single, fractionally small peripheral blood (PB) samples (3, 4). CTCs are extremely rare, and fewer than 1 CTC per mL of PB is associated with reduced overall survival for major cancers such as breast (5), colorectal (6), and prostate (7). Although there have been a number of large clinical studies in the last decade, the clinical value of CTC enumeration by liquid biopsy remains unclear (8–10). One major challenge is CTC heterogeneity, which has driven major efforts towards development “next-generation” liquid biopsy methods that permit genotypic and phenotypic characterization of single CTCs (11, 12).

A less-studied problem is that of *temporal* heterogeneity of CTCs and PB sampling, by which we mean the short term fluctuations in CTC numbers in PB that may be invisible to liquid biopsy. Liquid biopsy implicitly assumes that the number of CTCs in a blood sample is representative of the entire PB volume (PBV). Previous work has shown that this assumption may be statistically dubious in light of the rarity of CTCs and the fractionally small volume of samples (13, 14). With respect to the latter, the only currently FDA-cleared method for CTC enumeration in humans is CellSearch. In CellSearch 7.5 mL PB samples are analyzed which corresponds to about 0.015% of the ~5 L human PBV (6). Other experimental microfluidic platforms analyze similarly small samples in the range of 2-10 mL (0.004-0.02% PBV) (15–17). In pre-clinical mouse studies PB collection is limited to 200 μL every two weeks for non-terminal experiments (without fluid replacement), which is equivalent to about 10% of the ~1.5-2 mL mouse PBV (18, 19).

The small number of previously-published theoretical treatments of this problem assume that CTC detection statistics should follow a Poisson distribution (13, 20, 21). This implicitly assumes that CTCs are well-mixed in blood, and that the average number of CTCs in circulation does not change significantly over the minutes or hours surrounding the blood draw. Surprisingly, there is relatively little experimental pre-clinical or clinical data to support these assumptions (7, 20, 22, 23). The rarity of CTCs in mouse models often means that the entire peripheral blood volume must be drawn and analyzed, which is a terminal experiment that precludes serial study in the same mouse. Hence little is known about short-term fluctuations in CTC numbers *in vivo*.

‘*In vivo* flow cytometry’ (IVFC) is a general term for optical instrumentation designed to detect and enumerate circulating cells directly *in vivo*, most often using either fluorescence or photoacoustic contrast (19, 24). We recently developed ‘diffuse *in vivo* flow cytometry’ (DiFC) specifically for enumeration of rare green fluorescent protein (GFP)-expressing CTCs in mouse models of metastasis (25–29). DIFC uses diffuse light to non-invasively and continuously interrogate PB flowing in large, deeply-seated vessels (19). We recently used DiFC to monitor CTC dissemination in multiple myeloma (MM) and Lewis lung carcinoma (LLC) (27, 29) mouse xenograft models. We showed that DiFC permitted longitudinal study of CTCs and CTC multi-cellular clusters (CTCCs) in individual mice at burdens below 1 CTC per mL of PB.

In our previous work, we used DiFC only to study the *time-averaged* (mean) number of CTCs in circulation on a given day, but did not consider the *short-term* dynamics of CTC detections over timescales of minutes or hours. These short-term variations are recorded by DiFC, and therefore provide unique insights into temporal dynamics of CTCs *in vivo*. In this work, we analyzed DiFC data sets measured in mice and considered sample intervals equivalent to approximately 1%, 5%, 10% and 20% of the PBV. We analyzed how CTC numbers fluctuated over the timescales of minutes and hours. As we show, CTC numbers were far from steady state, and exhibited variability exceeding that expected by Poisson statistics and DiFC operator variability. This suggests that CTCs are not well-mixed in PB and that the mean number of CTCs changes over relatively short timescales. This can be explained by the short half-life of CTCs in circulation coupled with intermittent shedding of CTCs. These data provide direct *in vivo* evidence that CTC enumeration with fractionally small blood samples may fail to detect rare CTCs, and is generally quantitatively inaccurate. However, accuracy may be markedly improved by analysis of larger blood volumes, averaging of multiple blood samples, or continuous *in vivo* monitoring.

## 2. Materials and Methods

### 2.1 DiFC Instrument and Signal Processing

The DiFC instrument (***figure 1a***) and data processing algorithms were described in detail previously (26, 27, 29). Briefly, two specially designed fiber-optic probes (***fig. 1b***) are placed in-line along a major blood vessel (in this case, the ventral caudal bundle in the tail) as shown in ***fig. 1c***. The probes have integrated lenses and filters that allow efficient fluorescent light collection and rejection of non-specific tissue autofluorescence. GFP-expressing CTCs are detected by laser-induced fluorescence as they pass through the DiFC field of view (***fig. 1d***, also see ***supplementary methods S.1***). DiFC therefore permits detection of moving CTCs in large blood vessels 1-2 mm deep in tissue. As in our previous work, analysis of peak amplitude, width, and order of detection between the two channels allows us to discriminate arterial from venous flow directions (26). Detections of CTCs during a DiFC scan may be visualized using temporal raster plots, where each vertical black line represents a detection of a CTC (***fig. 1e***).

**Figure 1.**
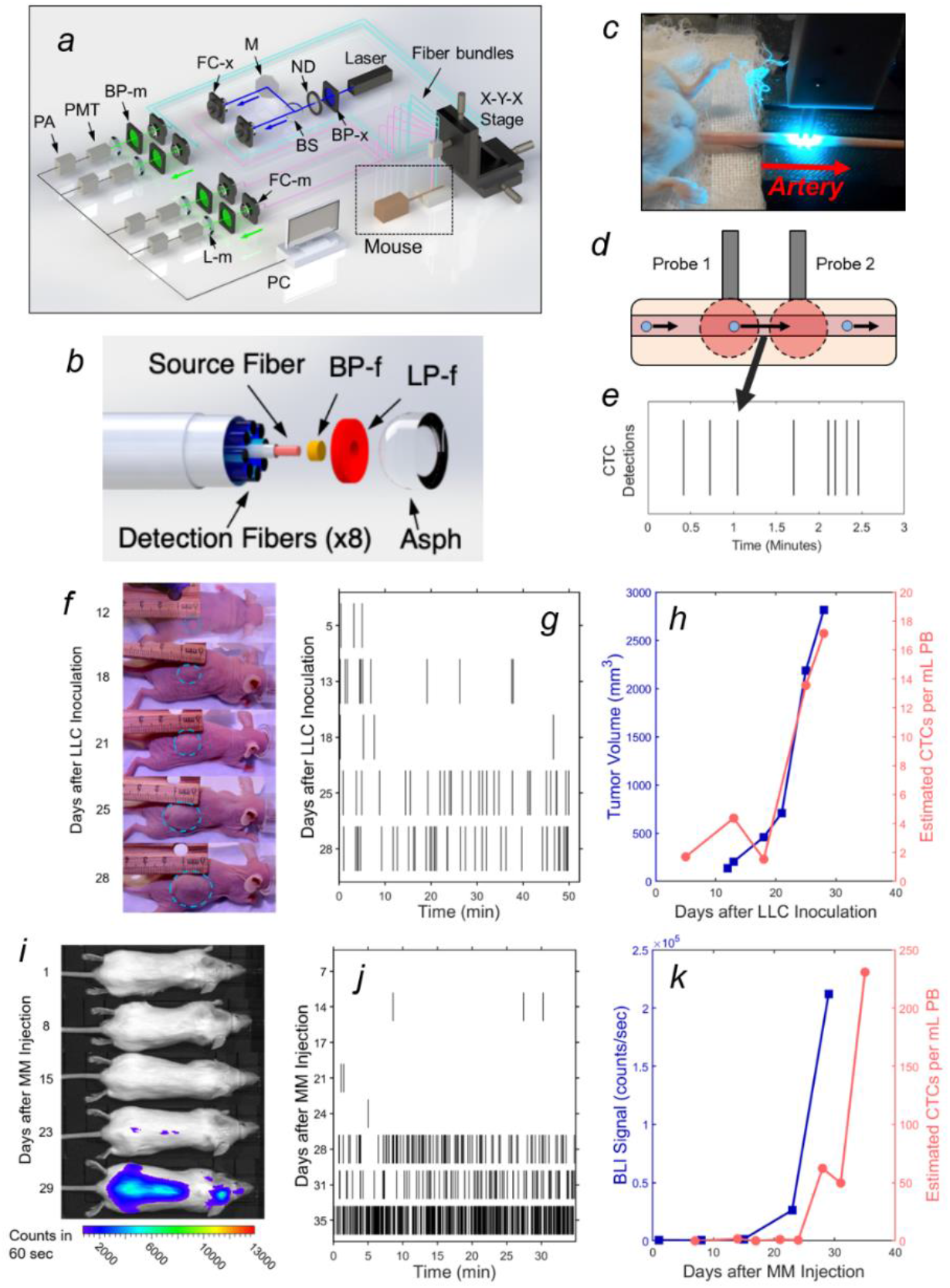
Diffuse in vivo flow cytometry (DiFC) is a system used to detect circulating tumor cells (CTCs) in mice. (a) The DiFC instrument was described in detail previously (27, 29). The specially designed DiFC probe uses two specially designed (b) optical fiber bundles that are placed on the skin surface approximately over a large vascular bundle, in this case (c) on the ventral side of the tail of a mouse. (d) When GFP-expressing CTCs pass through the DiFC field of view, the GFP is excited and the resulting fluorescent emission is detected by the fiber optic probes. (e) The emission is identified as a CTC detection in time, which is visualized as a vertical black line on a raster plot. (f-h) A representative data set collected from an s.c. LLC-tumor bearing mouse. (f) Photographs of LLC tumor growth are shown. (g) DiFC raster plots showing CTC detections during tumor grown, (h) The tumor volume (blue squares) and the mean rate of CTC detections (pink circles) over time is shown. (i-k) Representative DiFC data set collected from an MM-DXM mouse. (i) Bioluminescence (BLI) images during MM growth are shown. (j) DiFC raster plots of MM-CTC detections. (k) Plot of BLI signal (blue squares) and the mean rate of CTC detections (pink circles) over time. Figures (a) and (b) reproduced with permission from Patil et. al. (27).

We previously conservatively calculated that DiFC samples approximately 50 μL of PB per minute (27). As such, DiFC can sample the complete ~1.5-2 mL mouse PB volume (potentially more than once) in a single scan. Central to our analysis here is the assumption that the CTC count in a specific DiFC *time interval* is equivalent to the number of CTCs that would be counted in a *blood sample* of corresponding volume drawn from the same blood vessel at that time. For example, a 24 s DiFC scan interval is equivalent to approximately 20 μL of blood, or about 1% of the PBV of a mouse. In this work, we considered all 24 s, 2 min, 4 min and 8 min DiFC scan intervals, equivalent to 20, 100, 200 and 400 μL, or approximately 1%, 5%, 10% and 20% of the mouse PBV.

We characterized DiFC data sets in terms of the mean (μ) and variance (σ^2^) of CTC detections. In doing so we, analyzed all possible intervals in DiFC scans (a “sliding window”). While this approach yielded significant overlap (non-independence) between measurements, we showed explicitly that when large numbers of overlapping intervals are considered the variance converges to the non-overlapping case (see ***supplementary methods S.2***).

### 2.2 DiFC Data Sets

In this work, we analyzed five data sets as follows:

#### i) Lewis lung carcinoma (LLC) metastasis model

We re-analyzed previously reported DiFC data measured in sub-cutaneous (*s.c*.) Lewis lung carcinoma tumor bearing mice (29). As noted above, we previously considered only the mean CTC numbers on each day but not short-term fluctuations as we do here. Briefly, 10^6^ LLC cells expressing GFP (LL/2.GFP.Luc) cells were injected *s.c*. in the rear flank of nude mice and allowed to grow for up to 3 weeks. DiFC was performed for 40-50 minutes, at least once per week. CTCs were observed in circulation as early as 5 days after inoculation, and there was a general increase in CTC detection rate with increasing primary tumor volume (***figures 1f-h***). However, significant inter-mouse heterogeneity was observed in terms of both CTC numbers and lung metastasis formation. We used DiFC scans from this study where at least one CTC was detected (102 DiFC scans). CTC detection rates ranged from 0.019 to 1.05 CTCs per min, which is equivalent to 0.38 to 21.1 CTCs per mL of PB based on the DiFC sampling rate. We subsequently refer to these data as the “*LLC dataset*” below. A subset of 18 representative raster plots from this data set are shown in ***supplementary figure S1***.

#### ii) Multiple myeloma (MM) disseminated xenograft model (DXM)

We re-analyzed our previously reported DiFC data from an MM disseminated xenograft mouse model (DXM) (27). Briefly, 5 x 10^6^ GFP-expressing MM.1S.GFP.Luc cells were injected intravenously (*i.v.*) via the tail vein of SCID mice. After injection MM cells rapidly homed to the bone marrow niche, wherein they steadily proliferated and eventually re-entered circulation by the third week. DiFC was performed for 35 minutes twice weekly for up to 5 weeks. CTCs were relatively abundant (compared to the LLC model) and increased monotonically with bulk MM growth over time. Example data for an MM-DXM mouse is shown in ***figs. 1i-k***. In the analysis here, we used data sets from this study where DiFC detection rates exceeded 0.5 CTCs per minute only, and ranged from 0.6 to 19.6 CTCs per minute, which is equivalent to 12 to 392 CTCs per mL of PB. We refer to this as the “*MM 35-min dataset*” below. The complete set of DiFC raster plots are shown in ***supplementary figure S2***.

#### iii) 24-hour DiFC measurements in MM-DXM mice

We also performed new experiments in MM-DXM mice. All mice were handled in accordance with Northeastern University’s Institutional Animal Care and Use Committee (IACUC) policies on animal care. Animal experiments were carried out under Northeastern University IACUC protocol #15-0728R. All experiments and methods were performed with an approval from and in accordance with relevant guidelines and regulations of Northeastern University IACUC.

We performed *i.v*. injection of 5 x 10^6^ MM.1S.GFP.Luc cells in six, 8-week-old male severe combined immunodeficient SCID/Bg mice (Charles River) as in the MM-DXM model above (27). We performed cycles of four, 50-minute DiFC scans over a 24-hour period, beginning 4 weeks after inoculation. Institutional mouse housing followed a 7 AM-to-7 PM light, and 7 PM-to-7 AM dark cycle. To minimize possible bias in the start time for each cycle, the first DiFC scan was performed at either 7am (N = 7 data sets) or 7pm (N = 7 data sets). DiFC scans were performed by one of three human operators, and the first operator was also randomized. CTC detection rates for these data ranged from 0 to 42 CTCs per minute, equivalent to 0 to 840 CTCs per mL of blood. We refer to these as the “*MM 24-hour dataset*” below. The complete set of 24-hour DiFC measurements are shown in ***supplementary figure S3***.

Since 24-hour measurements required 3 human operators performing 4 alignments (physical repositioning) of the DiFC probe on the mouse tail in 24-hour periods, we also measured the inherent inter- and intra-operator variability in DiFC measurements (for details see ***supplementary methods S.3***). These data are referred to as “*1-operator-with-reposition dataset*” (N = 7 data sets) and “*2-operators-with-reposition dataset*” (N = 6 data sets) below.

#### iv) Limb-mimicking optical phantom with fluorescent microspheres

We also used a limb-mimicking optical flow phantom model as we have previously (27). We used Dragon Green fluorescence level 5 (DG5) reference microspheres (Cat. DG06M, Bangs Laboratories, Inc., Fishers, IN) suspended in PBS at concentrations of 200 or 400 microspheres per mL pumped through the phantom at linear flow speeds of 15, 30 or 60 mm/s. We performed DiFC at the 6 concentration-speed combinations in triplicate for 35 minutes in each case (N = 18). Microsphere suspensions were first sonicated to prevent aggregation and clumping and then mixed well to produce as homogenous a suspension as possible. In principle, this should yield DiFC detections that follow Poisson distributions. More details on these experiments are provided in ***supplementary methods S.4***.

#### v) Simulated DiFC data

We also simulated Poisson-distributed sequences of DiFC data *in silico*. We generated 54, 35-minute data sets using custom written code in Matlab (The Mathworks Inc., Natick, MA) with mean rates of detection in the same range as those measured in MM DXM mice. To compare MM mouse data to processes that are not described by a single Poisson, we also generated 35-minute Poisson simulations where the mean CTC detection rate increased by a factor of two halfway through the scan, i.e. Poisson detections with mean rate *λ*_1_ in the first 17.5 minutes and *λ*_2_ = 2*λ*_1_ in the second 17.5 minutes. We generated 54 such data sets. We also simulated DiFC data sets of the sum of two concurrent (simultaneous or merged) 35-minute, Poisson-distributed sequences of mean *λ*_1_ and *λ*_2_ = 2*λ*_1_ More details are provided in ***supplementary methods S.5***.

## 3. Results

### 3.1 For rare CTCs small samples frequently resulted in no CTC detections

We previously used DiFC to measure CTC dissemination in an LLC-tumor bearing mouse model (29), where CTCs were consistently rare throughout tumor growth. Specifically, in 102 DiFC scans, the CTC count rates corresponded to approximately 0.38 to 21.1 CTCs per mL of PB. Representative DiFC raster plots illustrating the range of the data set are shown in ***figures 2a-c***. Eighteen additional example data sets are also shown in ***figure S1***.

**Figure 2.**
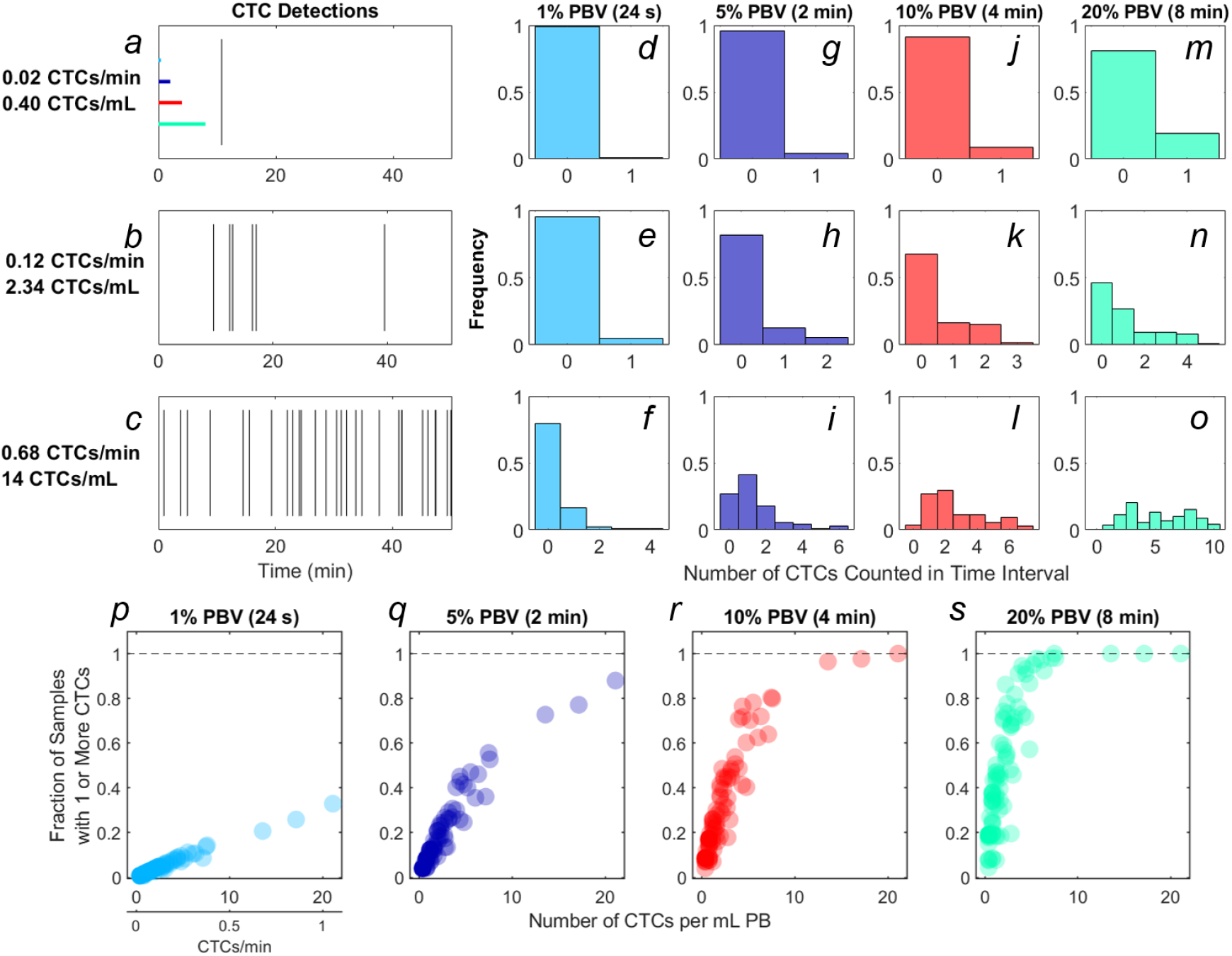
For rare CTCs, small blood samples frequently resulted in no DiFC detections. (a-c) Representative DiFC raster plots of LLC-CTC detections for average count rates of (a) 0.4, (b) 2.3 and (c) 14 CTCs per mL PB. Colored horizontal lines show the relative length of approximately 1% (24 s), 5% (2 min), 10% (4 min), and 20% (8 min) intervals (blood sample size). Distributions of CTC counts in detections intervals are shown for (d-f) 1% PBV, (g-i) 5%, (a4-c4) 10%, and (j-l) 20% PBV DiFC scan intervals. The fraction of intervals where 1 or more CTCs were detected for (p) 1%, (q) 5%, (r) 10%, and (s) 20% PBV scan intervals.

We considered the number of CTCs detected in 24 s, 2 min, 4 min and 8 min intervals during the DiFC scans. As above, these intervals were equivalent to approximately 20, 100, 200, and 400 μL of PB, or ~1%, 5%, 10% and 20% of the mouse PBV. Histograms of the number of CTC detections (for the data shown in ***figs. 2a-c***) are shown for 24 s (***figs. 2d-f***), 2 min (***figs. 2g-i***), 4 min (***fig. 2j-l***) and 8 min (***figs. 2m-o***) intervals, respectively. These data show that the probability that at least one CTC was detected in a small temporal sample was in general very low. For example, considering a 1% PBV sample size and 0.4 CTCs per mL, no CTCs were detected in 99% of equivalent blood samples (intervals) over the entire scan. Even for relatively high CTC burdens (14 CTCs per mL; ***fig. 2c***), 1% PBV sample sizes yielded zero CTC detections 79% of the time (***fig. 2f***). As would be expected, this probability improved significantly when larger time intervals (equivalent blood samples) were considered. For example, for a CTC burden of 14 CTCs per mL and an equivalent 20% PBV sample (***fig. 2o***), at least one CTC was detected 100% of the time.

Considering the data another way, the fraction of intervals for which at least 1 CTC was detected for all DiFC scans in the LLC dataset are shown in ***figures 2p-s***. In combination, these data underscore the fact that analysis of fractionally small blood samples (1-5% PBV) frequently resulted in detection of no CTCs, even though CTCs were present in the blood in all cases. Hence these data provide direct experimental validation of the notion that ‘more blood is better’ for detection of CTCs (14). Analysis of larger blood samples would further improve this: for example, analysis of 30% of the PBV would usually (> 50% of possible PBV samples) result in detection of at least 1 CTC in half of the LLC data sets here.

### 3.2 CTC counts in small samples were generally quantitively inaccurate

We also considered CTC enumeration accuracy from fractionally small blood samples. To study this, we used DiFC data measured in MM xenograft mice (“*MM 35-min dataset*”), (27) where CTCs were more abundant than in the LLC tumor model above. A representative 35-minute DiFC raster plot from an MM xenograft mouse, measured 31 days after engraftment of MM cells is shown in ***figure 3a***. (the full set of 18 DiFC scans from this data set are shown in ***figure S2***) The number of CTCs counted in sliding ~1%, 5%, 10% and 20% PBV intervals are shown in ***figs. 3b-e***, respectively. We also calculated the mean number of CTCs over the entire DiFC scan (black horizontal line in each case). By inspection, CTC counts varied significantly during the scan, with periods of relatively high and low detection rates.

**Figure 3.**
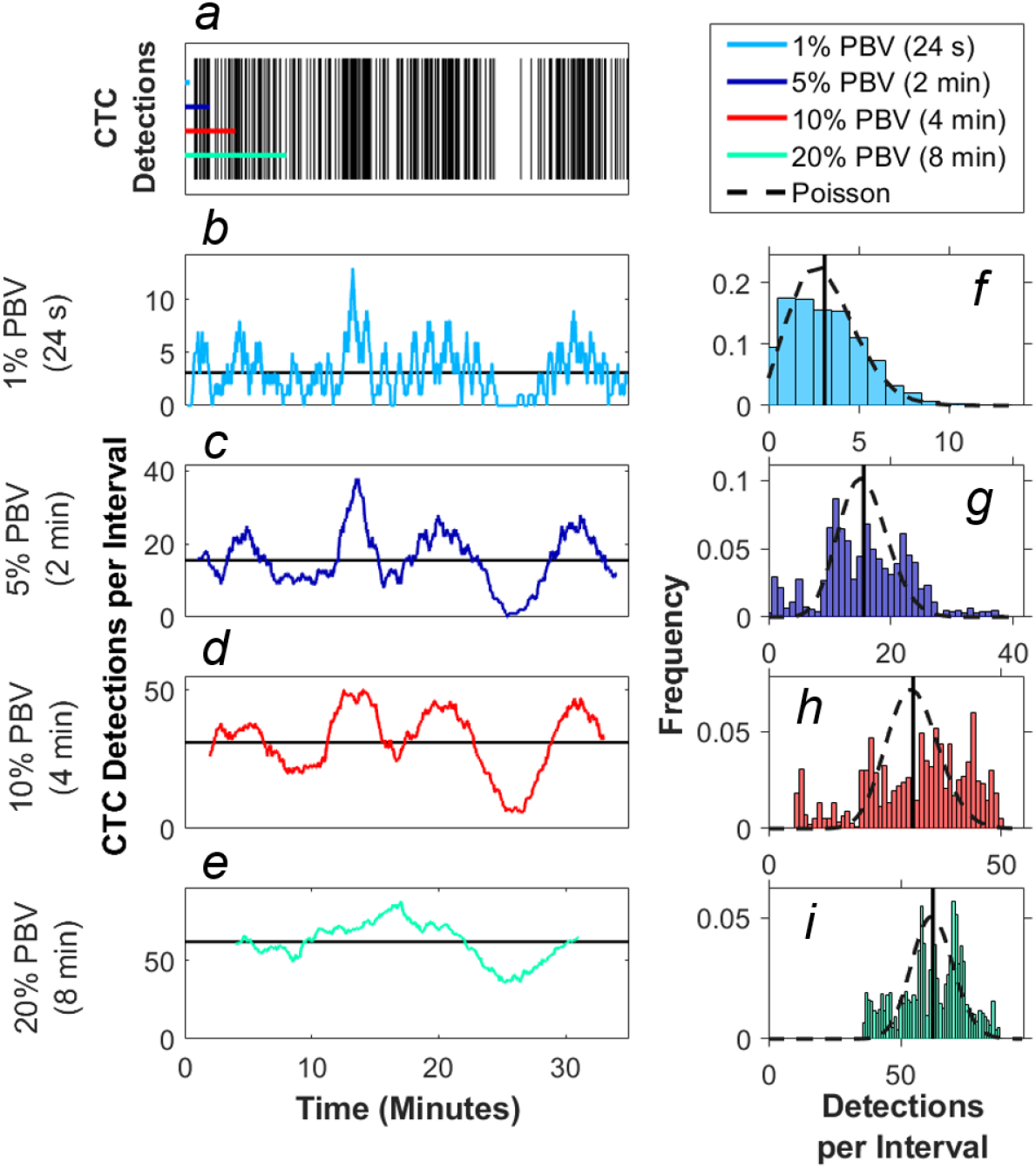
Significant fluctuation in CTC count rates were observed over the timescale of minutes in mice. (a) Representative DiFC raster plot measured from a MM DXM bearing mouse. Colored horizontal lines show the length of each time interval (blood sample size). The number of CTCs counted in sliding (b) 1%, (c) 5%, (d) 10% and (e) 20% equivalent PBV intervals are shown. Black horizontal lines identify the scan mean number of CTCs per interval. (f-i) The corresponding distributions of CTC counts for each equivalent PBV are shown. Black dashed lines indicate the distributions that would be expected from Poisson statistics. Black vertical lines show the mean number of CTCs.

These data demonstrate the surprisingly large range of CTC detection rates measured over 35-minute scans. For example, considering a 5% PBV interval (which is typical volume for a mouse blood collection experiment), equivalent detection rates ranged from 0 to 38 CTCs per sample. In other words, if PB was collected from this blood vessel, 100 μL samples drawn at different times (separated by just a few minutes) would have yielded vastly different (order-of-magnitude or more difference) numbers of CTCs.

The histograms of these data (i.e. the number of CTC detections for all 1%, 5%, 10% and 20% PBV intervals) are shown in ***figs. 3f-i***, along with the mean number of CTCs detected over the full scan (vertical black lines). We compared the measured distributions to Poisson distributions which, as noted above, are frequently assumed for liquid biopsy of PB (13, 20, 21). These are indicated by the black dotted curves in ***figs 3f-i***. By inspection, the DiFC-measured distribution of count rates did not appear to follow the Poisson distribution, particularly for larger (2-8 min) equivalent blood samples. The implications of this are discussed in more detail in section 3.3 below.

We also computed the ‘deviation from the scan mean’ (DFSM) for each observation (blood sample):

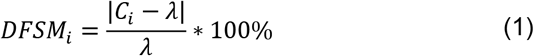

where *C_i_* is the number of CTCs in the *i^th^* equivalent sample, and *λ* is the mean number of CTC detections over the full scan for each sample size. The cumulative fraction of blood samples within 0-100% DFSM for the data in ***figs. 3f-i*** are shown in ***figure 4a***. The cumulative fraction of samples for Poisson distributions of the same mean are also shown (dotted lines, same colors), again suggesting that experimental data diverged substantially from the Poisson behavior.

**Figure 4.**
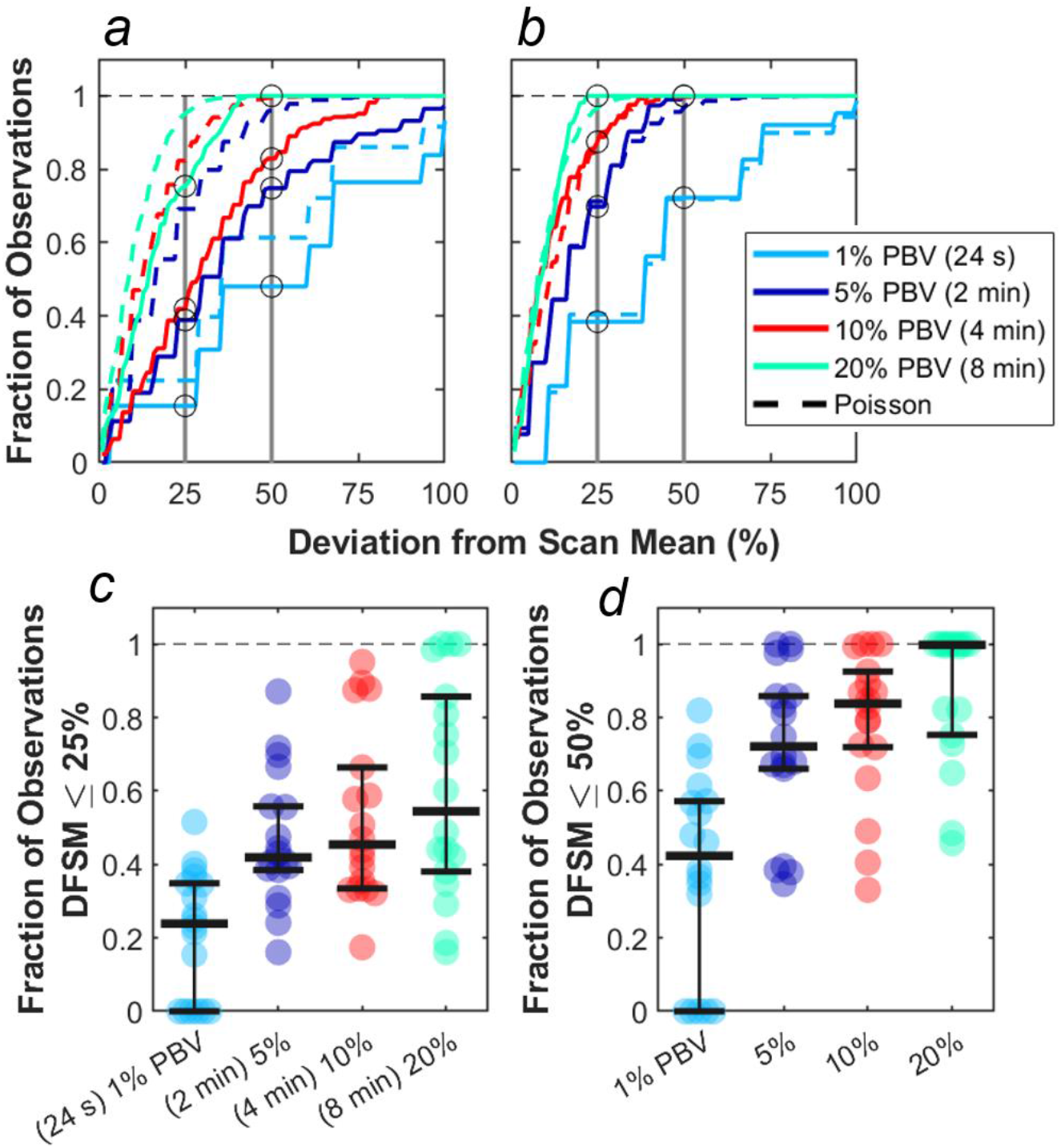
CTC counts in small equivalent blood samples were generally quantitatively inaccurate. (a) Analysis of a representative DiFC data set measured in an MM xenograft mouse, showing the impact of the sample size on the percentage deviation of CTC counts in an interval from the scan mean (DFSM; see text for details). The fraction of observations within a given DFSM are shown in each case. Dotted lines indicate the expected distribution based on Poisson statistics. (b) A second example data set that more closely followed the expected Poisson behavior. (c) The fraction of equivalent blood samples that were within 25% of the scan mean for all 18 DiFC scans in the MM 35-min dataset. (d) The fraction of samples within 50% of scan mean. Horizontal bars indicate the median, and first and third quartile for each blood sample size.

For example, these data can be interpreted as, “39% of randomly selected 5% PBV blood samples would yield a CTC count within 25% of the scan mean, whereas Poisson statistics predict that this number should be 69%”. It should also be noted that the large disagreement between expected Poisson behavior and measured behavior occurred in about half of the data sets. A representative plot from another DiFC scan where closer agreement was observed is shown in ***fig. 4b***.

We next considered the fraction of intervals where the DFSM was equal to or less than 25% and 50% for all sample sizes in the complete “*MM 35-min dataset*”. These data are summarized in ***figs. 4c,d***. Our use of 25% and 50% DFSM thresholds were selected since they are illustrative of sufficiently large error to affect prognostic classification – for example in determining whether a sample has 4 or 5 CTCs (breast and prostate cancer) or 2 or 3 CTCs (colorectal cancer) (6–8).

Taken together, these data show that quantitative estimation of CTC numbers from single samples in mice is extremely challenging. For example, for 5% PBV samples, the median probability for all scans of randomly obtaining a CTC count within 25% of the scan mean was only 41.9% (***fig. 4c***). Likewise, the probability of obtaining a CTC count within 50% of the mean was 72.1% (***fig. 4d***). All else being equal, use of larger blood sample volumes yielded higher probability of obtaining accurate count than smaller samples (14).

### 3.3 There were significant variations in CTC detection rates in 24 hour periods

We also studied the 24-hour variability in CTC detection rates in MM xenograft mice by performing four, 50 minute DiFC scans over 24-hour periods (“*MM 24-hour dataset*”). Half of the data sets began at 7PM, and half began at 7AM to rule out the possibility that the start time could affect the measurements. In addition, the starting DiFC human operator (one of 3) was randomized. Two representative DiFC data sets from 24-hour sessions are shown in ***figure 5*** with CTC detection rates of 2 min (5% PBV) intervals in ***figs. 5a,b***. The 50-minute average of each DiFC scan is shown (dotted horizontal lines), as well as the average over the 24-hour period (solid horizontal lines). The mean count rates over the 4 DiFC scans for both mice are summarized for clarity in ***fig. 5c***. As shown, the average measured detection rates changed by more than an order of magnitude depending on the time of day. The complete set of 14, 24-hour DiFC scans from this data set are shown in ***figure S3***.

**Figure 5.**
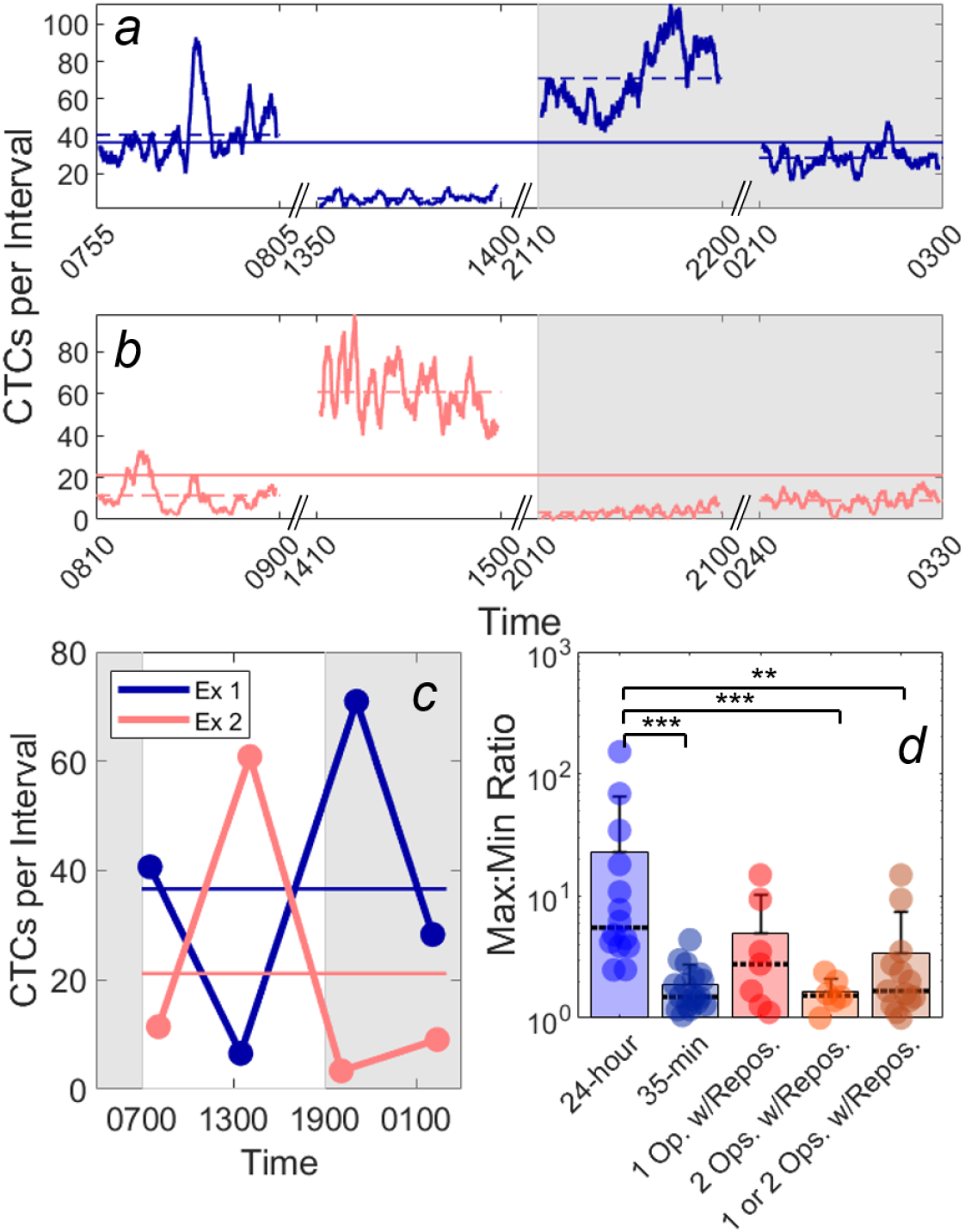
Large variations in DiFC count rate were observed over 24-hour periods. (a,b) Example DiFC measurements taken over 24 hours in two MM-bearing mice. The number of CTCs counted in 2 min (5% PBV) sliding intervals are shown. The solid horizontal lines in each figure show the 24-hour mean number of CTCs per 2 min interval, and the dashed horizontal lines show the local mean of the 50-minute DiFC scan. Dark hours are shaded grey. (c) The mean DiFC count rates over the 24-hour period is shown. (d) Ratios of maximum-to-minimum DiFC count rates are shown for MM 24-hour, MM 35-min (no repositioning), 1-operator-with-reposition, and 2-operators-with-reposition data sets. Bars represent the mean values with standard deviation error bars and dotted lines to identify the median values. Note the logarithmic y-axis.

To better quantify the 24-hour variability, the ratios of the maximum-to-minimum CTC detection rates for each scan are shown in the first column of ***fig. 5d***. As shown, this varied from 2.5 to 152. We note that we did not observe any specific circadian pattern to the data as others have reported for MM (23). Rather, the CTC count rates were seemingly randomly higher or lower at different times of days.

This 24-hour variability was also greater than the fluctuations observed over a short time scale (as in ***figures 3, 4***). To show this, DiFC scans from the “*MM 35-min dataset*” were each divided into the first 15 and last 15 minutes (separated by 5 minutes), and the same maximum-to-minimum count rates ratios were calculated. These ranged from 1.05 to 4.4 (***fig. 5d*** column 2). To rule out the possibility that the variability was due to repositioning of the DiFC probe on the skin surface between scans (which could affect the collection efficiency of the system), we tested the intra-operator reproducibility of DiFC count rates (“*1-operator-with-reposition dataset”*), and the corresponding maximum-to-minimum DiFC count rate ratios ranged from 1.1 to 14.9 (***fig. 5d*** column 3). We also tested the inter-operator reproducibility (“*2-operators-with-reposition dataset”*) of DiFC count rates, and the corresponding maximum-to-minimum ratios ranged from 1.0 to 2.4 (***fig. 5d*** column 4). When comparing *MM 24-hour* data to *MM 35-min, 1-operator-with-reposition, 2-operators-with-reposition*, and all reposition data by Kolmogorov-Smirnov test, p-values were p < 0.001, p = 0.058, p < 0.001, and p < 0.01 respectively. However, the intra- and inter-operator variability, as well as short-term fluctuations were not significantly different from each other (all p > 0.15). In summary, the variability in mean DiFC count rate measured over 24-hours was much larger than expected from repositioning variability.

### 3.4 Variability in CTC detection rates *in vivo* was higher than predicted by Poisson statistics

Because CTC detection is a random process some inherent measurement variability is expected. However, as already noted the variability observed in DiFC data *in vivo* frequently exceeded that expected by Poisson statistics. To further show this, we plotted the variance of the CTC counts per interval against the mean. This was repeated for 24 s, 2 min, 4 min and 8 min intervals. Since the mean and variance of an ideal Poisson distribution are equal, if Poisson distributed CTC measurements should fall approximately along the 1:1 curve. However, as shown in ***figures 6a-d***, the measurement variance for the “*MM 35-min dataset*” was larger than the mean for about half the measurements. This deviation was significantly larger when considering the “*MM 24-hour data sets*”, where fluctuations were observed over 24 hour periods (***figs. 6e-h***). Linear fits with a fixed intercept at zero resulted in slopes larger than 1, demonstrating that the variance generally exceed the mean.

**Figure 6.**
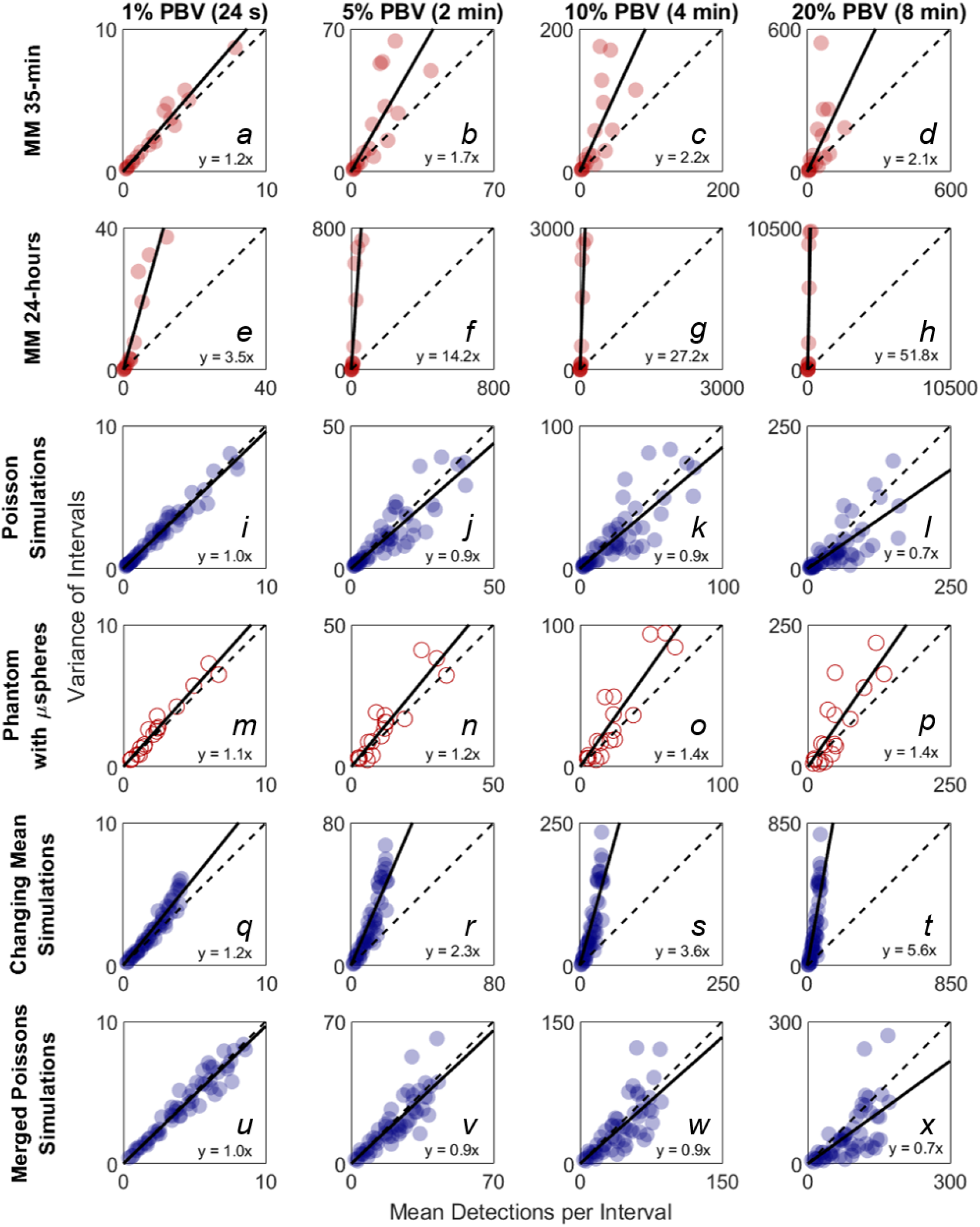
The variability in DiFC count rate measured in mice exceeded that expected by Poisson statistics. Measured variance in CTC counts compared to the scan mean count in DiFC data is shown for (a-d) MM 35-min mice, (e-h) MM 24-hour mice (i-l) simulated Poisson DiFC data, and (m-p) phantoms with suspensions of well-mixed microspheres. The dashed diagonal lines indicate the expected 1:1 variance-to-mean relationship for Poisson-distributed data. (q-t) Simulations show that the higher than expected variance in MM 35-min data is consistent with a Poisson process with changing mean (see text for details). (u-x) By contrast, two or more simultaneous (merged) Poisson processes would not be consistent with the higher-than-expected variance. Equations and solid lines indicate a linear fit to each data set.

To rule out the possibility that this deviation from Poisson behavior was an artifact of DiFC measurement or data analysis methods, we repeated the analysis on *in silico* simulated data sets, where detections were Poisson distributed as shown in ***figs. 6i-l***. We also performed DiFC measurements in a limb-mimicking optical flow phantom with suspensions of fluorescent microspheres that were well-mixed as shown in ***figs. 6m-p***. In both cases, the variance more closely agreed with the scan mean, and linear fitting yielded slopes that were generally close to 1. The slightly larger slopes observed in the phantom data likely resulted from microspheres settling in the syringe pump over the course of the DiFC scans, i.e. the spheres were not perfectly mixed.

The physical interpretation of these data is that the mean number of CTCs in the peripheral blood fluctuated significantly (i.e. was not in quasi-steady-state) over the time-scale of the DiFC experiments. To further test this, we simulated Poisson-distributed data sets where the mean number of CTCs doubled halfway through the scan (***figs. 6q-t)***. These data more closely resemble the *in vivo* experimental data (***figs. 6a-h***). It is also worth noting that the *in vivo* DiFC data is *not* consistent with multiple *concurrent* Poisson processes. As shown in ***figs. 6u-x*** (and by the properties of Poisson statistics) this summation would exhibit Poisson behavior.

The implications of these data are discussed in more detail below. However, in practice, this means simply that the expected variability in CTC numbers in a single, randomly-drawn blood sample is actually much worse (at least in this mouse model) than would be expected by Poisson statistics as others have assumed (13, 20).

### 3.5 Quantification of CTCs can be improved by averaging multiple samples

Previous studies have shown that analysis of larger blood samples provides more accurate quantification of CTCs than smaller blood samples (14). The large temporal variability and deviation from Poisson behavior observed in our data further predicts that averaging multiple small blood samples over 24-hour periods should yield more quantitatively accurate estimates (versus the average) than a single, larger blood sample. As summarized in ***figure 7***, we considered a number of sample sizes and sample number combinations We computed the number of samples where the DFSM ≤ 25% for each case. These are shown for (***fig. 7a***) 1%, 2%, two 1%, 4%, four 1%, 20%, 80%, and four 20% samples. Here, multiple samples were selected at least 6 hours apart.

**Figure 7.**
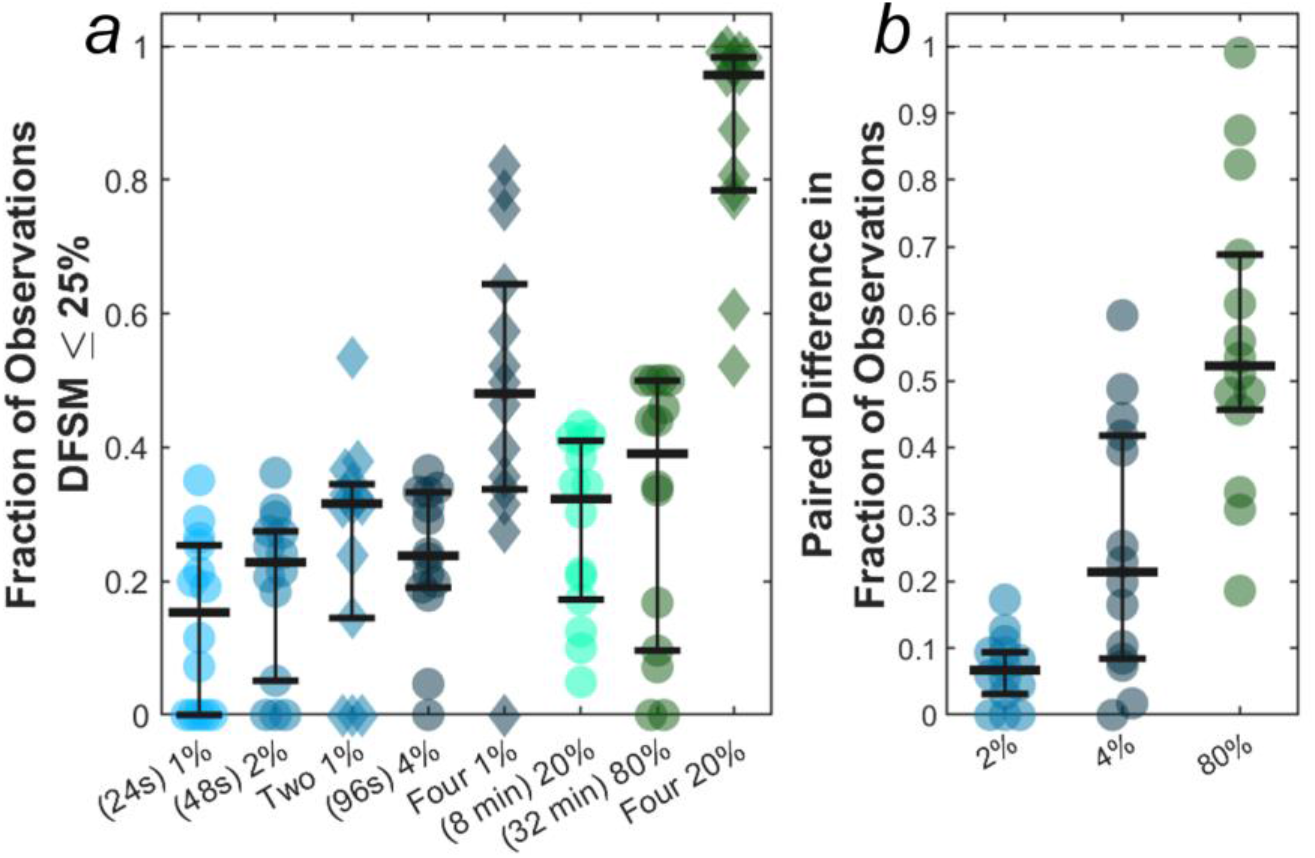
Use of larger blood samples and averaging multiple blood samples yields improved enumeration of CTCs. (a) Fraction of samples of different sizes where the DFSM was less than or equal to 25% for 1%, 2%, two 1%, 4%, four 1%, 20%, 80%, and four 20% PBV samples. (b) Paired difference in the fraction of samples were the DFSM <= 25% between the averaged samples (two 1%, four 1%, and four 20% PBV) and a single continuous sample of equivalent size (2%, 4%, and 80% PBV).

We then compared paired differences for equivalent total blood volumes taken continuously or throughout the 24-hour period as shown in ***fig. 7b***. For example, we compared two, averaged 1% PBV (24s) samples to a single 2% PBV (2 min) measurement. Similarly, we compared four, 1% PBV samples (each taken at least 6 hours apart) to a single 4% PBV sample, as well as four, 20% PBV samples to one single 80% PBV sample (32 min). For nearly every data set, averaging multiple smaller blood samples yielded a higher likelihood of accurately estimating the 24-hour CTC rate compared to single, larger blood samples. This data demonstrates that while larger blood samples do improve the likelihood of accurately estimating CTC numbers, further improvements may be made by averaging smaller samples taken at different times of the day.

## 4. Discussion

Although CTCs are widely studied using liquid biopsy, analysis of single, fractionally-small PB samples is inherently blind to temporal fluctuations in CTC numbers that occur over the timescale of minutes or hours. To our knowledge there is little clinical or pre-clinical evidence supporting the implicit assumption that CTC numbers in PB are relatively stable over the minutes and hours surrounding the blood draw. As we have noted, DiFC samples large volumes of circulating PB continuously and non-invasively, and therefore provides unique insights into CTC dynamics that occur *in vivo* in mouse models of metastasis. These data showed that for rare CTCs (LLC bearing mice), any single small sampling interval (equivalent blood sample) frequently resulted in no CTC detections. For more abundant CTCs (MM bearing mice), small samples were unlikely to yield a quantitatively accurate estimate of mean CTC numbers. Data taken over 24-hour periods also showed that CTC numbers varied by up to two orders-of-magnitude, with variance well in excess of that expected by Poisson statistics or operator variability.

As we have discussed, in combination, these imply that CTCs are not well mixed in the PB. This is consistent with two established properties of CTCs, i) that CTCs have a short half-life in circulation, and ii) that CTCs shed continuously from primary tumor(s). For the former, the small number of clinical estimates of CTC half-life range between 6 minutes and 2.4 hours (15, 30, 31). Estimates of CTC half-life in mouse models also range from 10 to 60 minutes for different cell types and mouse strains (29, 32–37). In addition, many CTCs are cleared in minutes in the “first-pass effect” in the lungs, liver, or spleen (38, 39). With respect to the kinetics of CTC shedding, it has been previously shown that tumors disseminate CTCs at a rate of about 10^6^ cells per gram of tumor tissue per day (40). We were unable to find any specific information on the short-term dynamics of CTC shedding from the primary tumor, however it is conceivable that this – like other tumor processes - varies over the timescale of minutes. For example, it has been shown that tumors cycle through hypoxia states on similar timescales (41).

These phenomena suggest that CTCs are continuously and dynamically being shed into and cleared from circulation. For example, in one LLC data set (***fig. 3a***), only a single detection was observed in a 50-minute DiFC scan suggesting that the CTC cleared from circulation before it could be detected a second time.

DiFC and simulation data in this model suggest that CTC numbers may be better described using a kinetic model that oscillates between states of relatively high and low shedding as opposed to a steady Poisson process.

Specifically, measured CTC data in MM-DXM bearing mice was more consistent with our *in silico* simulated data sets where the mean CTC rate increased partway through the scan (***figs. 6q-t***). These simulations suggest that CTC rates may change by approximately a factor of two over a 35-minute scan, and by larger factors over 24-hour periods. The disseminated nature of the MM tumor also implies that there may be multiple (many) sites of CTC shedding. However, as we noted, multiple simultaneous Poisson-distributed CTC sources would not produce the observed in vivo DiFC data (***figs. 6u-x***). This suggests that the changing numbers of CTCs in circulation is in response to systemic factors such as hormonal or cardiovascular effects. Therefore, even in a disseminated tumor model, there are further factors causing systemic CTC rate variations over time.

Despite the similarities in data, our simulations of a doubling in detection rate midway through the scan is, of course, not necessarily representative of what occurs biologically. In general, the magnitude and frequency of CTC rate changes is not known, and moreover is expected to vary with cancer type and mouse strain. Determining an accurate model of these changes is the subject of ongoing work in our group.

Since our observations were made in mouse xenograft models, a natural question is whether similar short-term temporal fluctuations occur in humans. The general findings here are consistent with the small number of published studies in the literature. Martin et. al. (22) studied CTC numbers in blood samples taken at 12-hour intervals from 51 breast cancer patients. While they concluded that there was no diurnal pattern, blood samples taken on the same day yielded differences by up to a factor of 3.8. Some patients had CTCs detected in one blood sample and no CTCs in the second blood sample, and several switched between prognostic categories (i.e. numbers above or below the 5 CTC threshold). Likewise, Paiva *et. al*. (23) showed 24-hour variability of up to a factor of 7 in MM patients, and suggested a circadian rhythm to the fluctuations. Juratli *et. al*. (42) took three sequential 3 mL PB samples from each of 7 stage IV melanoma patients, and found also found significant variation in CTC number (by up to a factor of 4), including four patients where no CTCs were detected in some of the samples.

Together our findings suggest a number of practical implications for the use of liquid biopsy in CTC enumeration. As noted the clinical value of CTC numbers as a diagnostic or predictive biomarker is still not clear (8–10, 43, 44), which has driven major efforts towards development of next-generation liquid biopsy technology for characterization of CTC genotypic and phenotypic heterogeneity (12–14). Assuming that the CTC behavior observed here extends to humans, this may provide a simple complementary explanation (to CTC heterogeneity) for these challenges: it is inherently difficult to enumerate CTCs from fractionally small blood samples because of the natural temporal fluctuations. As shown, these fluctuations could certainly result in change of prognostic category (CTC positive or negative) when simple numerical thresholds are applied.

Second, these data are consistent with the idea that improved clinical CTC enumeration could be achieved by analysis of larger blood samples (13, 14), either by liquid biopsy or by development of *in vivo* methods for counting or capturing CTCs (45). Our analysis suggests that averaging of multiple small blood samples taken over the course of hours should yield even further improved estimation of CTC numbers. Alternatively, new in vivo methods could scan larger blood volumes over time and in principle yield more accurate enumeration of CTCs. In this regard, because DiFC is inherently scalable to larger tissue volumes, we are already exploring the possible translation of DiFC to humans through the use of highly specific fluorescence molecular probes (46).

Third, CTC clusters (CTCCs) are even more rare than CTCs (they occur at a frequency of less than 10% of single CTCs) but are of great interest because they are known to have significantly higher metastatic potential (47). Although we did not explicitly consider CTCCs in the analysis here, DiFC does permit detection of CTCCs (27, 29). The relative rarity of CTCCs implies that the challenges of enumeration with liquid biopsy of small PB samples (i.e. the low probability of detection and poor likelihood of accurate quantification) are likely to be further exacerbated. This is also true when considering accurate enumeration of specific (and more-rare) CTC phenotypes (11).

Last, the fast turnover of CTCs in peripheral blood also suggests that anti-CTC therapeutic strategies such as “CTC dialysis” that have been proposed (48–50) are unlikely to succeed unless performed continuously, for example using a wearable device.

In summary, analysis of DiFC data in mouse models of metastasis shows that CTC numbers are far from steady-state *in vivo* and undergo significant fluctuations on the timescales of minutes and hours. This can cause significant error in CTC detection and enumeration using small blood samples and motivates new methods for analyzing larger blood volumes *in vivo*. Ongoing work by our team includes the application of DiFC to other mouse xenograft models, investigating future clinical use of DiFC, and development of mathematical models to more accurately describe CTC dynamics.

## Acknowledgments

This work was supported by the National Institutes of Health (NIH) (R01HL124315; NHLBI). We thank Dr. Paul Mathew (Tufts Medical Center, Boston, MA), Prof. Dana Brooks (Northeastern University, Boston, MA), and Prof. Daniel Ocone (Rutgers University, Piscataway, NJ) for their helpful comments and suggestions on our work. We also sincerely thank Dr. Miguel Martin (Universidad Complutense, Madrid, Spain) for kindly sharing CTC data.

## Notes

### Competing Interest Statement

The authors have declared no competing interest.

### Summary of Updates

New sets of simulated data were performed to compare in vivo data to theoretical biological models of CTC dynamics. In vivo data is similar to simulations of Poisson processes with a changing mean, suggesting CTC shedding and/or clearance rates fluctuate over 35-min and 24-hour periods. Additionally, a new author has been added, figures 3-7 revised, and supplemental materials updated.

## References

1. G. P. Gupta, J. Massague, Cancer metastasis: building a framework. Cell 127, 679–695 (2006).

2. P. S. Steeg, D. Theodorescu, Metastasis: a therapeutic target for cancer. Nat Clin Pract Oncol 5, 206–219 (2008).

3. C. Alix-Panabieres, K. Pantel, Circulating tumor cells: liquid biopsy of cancer. Clin Chem 59, 110–118 (2013).

4. S. Mader, K. Pantel, Liquid biopsy: current status and future perspectives. Oncol Res Treat 40, 404–408 (2017).

5. M. Cristofanilli et al., Circulating tumor cells, disease progression, and survival in metastatic breast cancer. N Engl J Med 351, 781–791 (2004).

6. S. J. Cohen et al., Relationship of circulating tumor cells to tumor response, progression-free survival, and overall survival in patients with metastatic colorectal cancer. J Clin Oncol 26, 3213–3221 (2008).

7. J. G. Moreno et al., Circulating tumor cells predict survival in patients with metastatic prostate cancer. Urology 65, 713–718 (2005).

8. J. B. Smerage et al., Circulating tumor cells and response to chemotherapy in metastatic breast cancer: SWOG S0500. J Clin Oncol 32, 3483–3489 (2014).

9. C. Raimondi, A. Gradilone, G. Naso, E. Cortesi, P. Gazzaniga, Clinical utility of circulating tumor cell counting through CellSearch((R)): the dilemma of a concept suspended in Limbo. Onco Targets Ther 7, 619–625 (2014).

10. C. Alix-Panabieres, K. Pantel, Challenges in circulating tumour cell research. Nat Rev Cancer 14, 623–631 (2014).

11. B. Polzer et al., Molecular profiling of single circulating tumor cells with diagnostic intention. EMBO Mol Med 6, 1371–1386 (2014).

12. A. Bardia, D. A. Haber, Solidifying liquid biopsies: can circulating tumor cell monitoring guide treatment selection in breast cancer? J Clin Oncol 32, 3470–3471 (2014).

13. A. L. Allan, M. Keeney, Circulating tumor cell analysis: technical and statistical considerations for application to the clinic. J Oncol 2010, 426218 (2010).

14. Z. S. Lalmahomed et al., Circulating tumor cells and sample size: the more, the better. J Clin Oncol 28, e288–289; author reply e290 (2010).

15. S. L. Stott et al., Isolation and characterization of circulating tumor cells from patients with localized and metastatic prostate cancer. Sci Transl Med 2, 25ra23 (2010).

16. N. M. Karabacak et al., Microfluidic, marker-free isolation of circulating tumor cells from blood samples. Nat Protoc 9, 694–710 (2014).

17. S. H. Au et al., Microfluidic Isolation of Circulating Tumor Cell Clusters by Size and Asymmetry. Sci Rep 7, 2433 (2017).

18. J. Hoff, Methods of blood collection in the mouse. Lab Animal 29, 47–53 (2000).

19. C. Hartmann, R. Patil, C. P. Lin, M. Niedre, Fluorescence detection, enumeration and characterization of single circulating cells in vivo: technology, applications and future prospects. Phys Med Biol 63, 01TR01 (2017).

20. A. G. Tibbe, M. C. Miller, L. W. Terstappen, Statistical considerations for enumeration of circulating tumor cells. Cytometry A 71, 154–162 (2007).

21. F. A. Coumans, S. T. Ligthart, L. W. Terstappen, Interpretation of changes in circulating tumor cell counts. Transl Oncol 5, 486–491 (2012).

22. M. Martin et al., Circulating tumor cells in metastatic breast cancer: timing of blood extraction for analysis. Anticancer Res 29, 4185–4187 (2009).

23. B. Paiva et al., Detailed characterization of multiple myeloma circulating tumor cells shows unique phenotypic, cytogenetic, functional, and circadian distribution profile. Blood 122, 3591–3598 (2013).

24. V. V. Tuchin, A. Tarnok, V. P. Zharov, In vivo flow cytometry: a horizon of opportunities. Cytometry A 79, 737–745 (2011).

25. E. Zettergren et al., Instrument for fluorescence sensing of circulating cells with diffuse light in mice in vivo. J Biomed Opt 17, 037001 (2012).

26. X. Tan et al., In vivo flow cytometry of extremely rare circulating cells. Sci Rep 9, 3366 (2019).

27. R. Patil et al., Fluorescence monitoring of rare circulating tumor cell and cluster dissemination in a multiple myeloma xenograft model in vivo. J Biomed Opt 24, 1–11 (2019).

28. W. Di et al., Real-time particle-by-particle detection of erythrocyte-camouflaged microsensor with extended circulation time in the bloodstream. Proc Natl Acad Sci U S A 117, 3509–3517 (2020).

29. J. E. Fitzgerald et al., Heterogeneity of circulating tumor cell dissemination and lung metastases in a subcutaneous Lewis lung carcinoma model. Biomed Opt Express 11, 3633–3647 (2020).

30. S. Meng et al., Circulating tumor cells in patients with breast cancer dormancy. Clin Cancer Res 10, 8152–8162 (2004).

31. N. Aceto et al., Circulating tumor cell clusters are oligoclonal precursors of breast cancer metastasis. Cell 158, 1110–1122 (2014).

32. I. Georgakoudi et al., In vivo flow cytometry: a new method for enumerating circulating cancer cells. Cancer Res 64, 5044–5047 (2004).

33. D. A. Sipkins et al., In vivo imaging of specialized bone marrow endothelial microdomains for tumour engraftment. Nature 435, 969–973 (2005).

34. W. He, H. Wang, L. C. Hartmann, J. X. Cheng, P. S. Low, In vivo quantitation of rare circulating tumor cells by multiphoton intravital flow cytometry. Proc Natl Acad Sci U S A 104, 11760–11765 (2007).

35. J. M. Runnels et al., Optical techniques for tracking multiple myeloma engraftment, growth, and response to therapy. J Biomed Opt 16, 011006 (2011).

36. N. Pestana et al., Improved diffuse fluorescence flow cytometer prototype for high sensitivity detection of rare circulating cells in vivo. J Biomed Opt 18, 077002 (2013).

37. V. Pera et al., Diffuse fluorescence fiber probe for in vivo detection of circulating cells. J Biomed Opt 22, 37004 (2017).

38. N. Mizuno, Y. Kato, Y. Izumi, T. Irimura, Y. Sugiyama, Importance of hepatic first-pass removal in metastasis of colon carcinoma cells. J Hepatol 28, 865–877 (1998).

39. U. M. Fischer et al., Pulmonary passage is a major obstacle for intravenous stem cell delivery: the pulmonary first-pass effect. Stem Cells Dev 18, 683–692 (2009).

40. T. P. Butler, P. M. Gullino, Quantitation of cell shedding into efferent blood of mammary adenocarcinoma. Cancer Res 35, 512–516 (1975).

41. M. W. Dewhirst, Relationships between cycling hypoxia, HIF-1, angiogenesis and oxidative stress. Radiat Res 172, 653–665 (2009).

42. M. A. Juratli et al., Dynamic fluctuation of circulating tumor cells during cancer progression. Cancers (Basel) 6, 128–142 (2014).

43. J. S. de Bono et al., Circulating tumor cells predict survival benefit from treatment in metastatic castration-resistant prostate cancer. Clin Cancer Res 14, 6302–6309 (2008).

44. A. Toss, Z. Mu, S. Fernandez, M. Cristofanilli, CTC enumeration and characterization: moving toward personalized medicine. Ann Transl Med 2, 108 (2014).

45. N. Saucedo-Zeni et al., A novel method for the in vivo isolation of circulating tumor cells from peripheral blood of cancer patients using a functionalized and structured medical wire. Int J Oncol 41, 1241–1250 (2012).

46. R. A. Patil, M. Srinivasarao, M. M. Amiji, P. S. Low, M. Niedre, Fluorescence labeling of circulating tumor cells with a folate receptor-targeted molecular probe for diffuse In vivo flow cytometry. Mol Imaging Biol 10.1007/s11307-020-01505-9 (2020).

47. Y. Hong, F. Fang, Q. Zhang, Circulating tumor cell clusters: What we know and what we expect (Review). Int J Oncol 49, 2206–2216 (2016).

48. B. Faltas, Cornering metastases: therapeutic targeting of circulating tumor cells and stem cells. Front Oncol 2, 68 (2012).

49. Y. R. Kim, J. K. Yoo, C. W. Jeong, J. W. Choi, Selective killing of circulating tumor cells prevents metastasis and extends survival. J Hematol Oncol 11, 114 (2018).

50. E. I. Galanzha et al., In vivo liquid biopsy using Cytophone platform for photoacoustic detection of circulating tumor cells in patients with melanoma. Sci Transl Med 11 (2019).

