## Supplementary Materials for "Short-term circulating tumor cell dynamics in mouse xenograft models and implications for liquid biopsy"

ORCID: A.L.W: 0000-0003-1815-6479, J.E.F: 0000-0001-9694-5540, F.I.: 0000-0003-2830-3161

**This PDF file includes:**

Supplementary Methods
Figures S1 to S3

### Supplementary Methods

#### S.1 DiFC Instrument and Signal Processing

The DiFC instrument schematic is shown in ***figure 1a***, which was described in detail previously (1). It is described here in brief for completeness. The light source was a 488 nm DPSS laser, the output of which was filtered with a 488/10 nm cleanup band-pass (BP-x). The laser was coupled into the fiber probe central "source fiber" using lens-fiber couplers (FC-x) with 532 nm anti-reflection coating. The fiber probes were custom designed and built for this application by EMVision LLC (Loxahatchee, FL). Two filters were mounted directly to the tip of the fiber bundles, a central 488/5 nm band-pass filter (BP-f) for the excitation light, and an outer detection ring shape 503 nm long-pass filter (LP-f) for collection of fluorescence light. We found both to be necessary to mitigate fiber autofluorescence (***fig. 1b***). The outputs of the collection fiber bundles were terminated on a second set of lens fiber couplers (FC-m), and then filtered with two identical interference filters (BP-m) at 535/50 nm. Four photomultiplier tubes (PMT) were powered by stable PMT voltage supplies. The output of each PMT was amplified with low-noise pre-amplifiers with 300 Hz low pass filter, and then digitized with a multi-function data acquisition card. The entire setup was mounted on an optics cart so that it could be easily between sites. Data collection software was specially written in Matlab.

Data processing was performed as previously described (1) with one small exception. As discussed in Fitzgerald et. al. (2), PMTs were upgraded to lower noise versions after publication of Patil et. al. (1). As such, data processing of the new DiFC data sets in this paper (specifically: "*MM 24-hour*", "*1-operator-with-reposition*", "*2-operators-with-reposition*"), a detection threshold of 7 mV was used. This was empirically determined to yield a low false alarm rate and high sensitivity for the new detectors.

#### S.2 Proof That the Mean and Variance of Overlapping and Non-Overlapping DiFC Data Intervals Converge for Large Numbers of Intervals.

Throughout the manuscript we analyzed overlapping (sliding window) intervals of DiFC data for calculation of the variance of CTC arrival times. The number of CTC arrivals in two overlapping windows are not independent, which means that there are no guarantees that the sample variance should be unbiased estimator of the true variance. Nonetheless, we next show that this approach which has the advantage that it provides much higher numbers of intervals for our analysis, but yields the same variance as non-overlapping intervals, provided large (>100) numbers of intervals are considered. The proof is as follows.

Let $\{X_1, \ldots, X_n\}$ be a sequence of identically distributed random variables, with mean $\mu$ and variance $\sigma^2$. In the event estimation application, $X_i$ is the random variable that counts the number of events in the $i^{th}$ sliding window. Suppose that $m$ is a given integer, $m \ll n$, and that the each of following subsets of random variables is independent:

$$S_1 = \{X_1, X_{1+m}, X_{1+2m}, \ldots\}$$
$$S_2 = \{X_2, X_{2+m}, X_{2+2m}, \ldots\}$$
$$\vdots$$
$$S_{m-1} = \{X_{m-1}, X_{m-1+m}, X_{m-1+2m}, \ldots\}$$
$$S_m = \{X_m, X_{m+m}, X_{m+2m}, \ldots\}$$

In the event estimation application, $m$ would be the window size, and we assume that events in disjoint intervals are independent as they would be for a Poisson process.

For simplicity, assume that $n$ is a multiple of $m$, so that $r = n/m$ is an integer. The number of variables in each set is roughly $r$ (not exactly, since there is no "$X_{1+rm}$" in the first set, and so forth, but this a good approximation for large $n$).

Now we compute the sample variance

$$\bar{\sigma}^2 = \frac{1}{n-1}\sum_{i=1}^{n}(x_i - \bar{\mu})^2$$

from a sample $\{x_1, \dots, x_n\}$ drawn from $\{X_1, \dots, X_n\}$, where $\bar{\mu}$ is the sample mean. We claim that, for large $n$, $\bar{\sigma}^2 \approx \sigma^2$ and $\bar{\mu}^2 \approx \mu^2$.

As a corollary, if the variables $X_i$ are Poisson distributed, it will follow that $\bar{\sigma}^2 \approx \bar{\mu}$.

Because $n$ is large, we can approximate the sample variance by dividing by $n$ instead of $n-1$:

$$\bar{\sigma}^2 \approx \frac{1}{n}\sum_{i=1}^{n}(x_i - \bar{\mu})^2.$$

The sample mean $\bar{\mu} = \frac{1}{n}\sum_{i=1}^{n} x_i$ has expected value $\frac{1}{n}\sum_{i=1}^{n}\mu = \mu$ (this does not require independence) so, again because $n$ is large, by the law of large numbers we are justified in approximating $\bar{\mu} \approx \mu$.

In conclusion, we will assume that

$$\bar{\sigma}^2 \approx \frac{1}{n}\sum_{i=1}^{n}(x_i - \mu)^2 \approx \Sigma_1 + \cdots + \Sigma_m,$$

where we have split the sum according to the subsets $S_j$, $j = 1, \dots, m$, with respective partial sums:

$$\Sigma_j = \frac{1}{n}\sum_{i=0}^{r-1}\left(x_{j+im} - \mu\right)^2$$

(As mentioned earlier, we are truncating to avoid the few last terms that are not there).

Since the variables in each set $S_j$ are independent, these terms represent IID samples from the same distribution, and therefore the expected value of

$$\frac{n}{r}\Sigma_j = \frac{1}{n}\sum_{i=0}^{r-1}\left(x_{j+im} - \mu\right)^2$$

(approximating $r \approx r-1$) is the variance $\sigma^2$. Therefore

$$E\left[\Sigma_j\right] \approx \frac{r}{n}\sigma^2.$$

Since

$$E[\bar{\sigma}^2] \approx \sum_{i=1}^{m} E\left[\Sigma_j\right] \approx m\frac{r}{n}\sigma^2 = \sigma^2,$$

we conclude that $E[\bar{\sigma}^2] \approx \sigma^2$, so by the law of large numbers, $\bar{\sigma}^2 \approx \sigma^2$, as claimed.

#### S.3 Estimation of Variability in CTC Count Rate Due to Inter- and Intra- DiFC Operator Experimental Variability.

As noted in section 2.2, the "*MM 24-hour dataset*" required 3 human operators performing 4 alignments (physical repositioning) of the DiFC probe on the mouse tail over a 24-hour period. To account for the possibility that variations in CTC count rates were due to probe repositioning (as opposed to real differences in CTC numbers), we checked reproducibility of DiFC measurements in an additional set of MM-DXM mice as follows. Mice were injected intravenously with MM.1S cells as described in the main body of the text.

One operator positioned the DiFC probe and acquired 15 minutes of data, and then removed the probe from the mouse tail. The same operator repositioned the probe on the tail surface and acquired another 15 minutes of data. Due to process of repositioning, approximately 30 minutes transpired between the end of the first scan and the start of the second scan. These data are referred to as "*1-operator-with-reposition dataset*" (N = 7 data sets) in the text.

Second, one operator positioned the DiFC probe and acquired 15 minutes of data, and then removed the probe from the mouse tail. A different (second) operator repositioned the DiFC probe on the tail and acquired another 15 minutes of data. Again, about 30 minutes separated these two scans. These data set are referred to as "2-*operators-with-reposition dataset*" (N = 6 data sets) in the text.

### S.4 Description of the Limb-Mimicking Optical Flow Phantoms

As described in section 2.2, we used an optical flow phantom model with suspensions of fluorescent microspheres that were well-mixed, as we have done previously (1). The purpose of the experiment in this work is to produce an experimental case where DiFC detections should follow Poisson statistics. Specifically, we used Dragon Green fluorescence level 5 (DG5) reference standard microspheres (Cat. DG06M, Bangs Laboratories, Inc., Fishers, IN) which have similar size and fluorescence intensity as a bright GFP-expressing cell. Microspheres were suspended in PBS at concentrations of approximately 200 and 400 μspheres per mL and the suspensions were sonicated (Sonicator info here) to ensure even microsphere distribution. These suspensions were passed through an optical phantom made from high-density polyethylene (HDPE). We showed previously that this model has similar optical properties as biological tissue.

Tygon tubing (TGY-010-C, Small Parts, Inc., Seattle, WA) was threaded through a 1 mm diameter channel in the phantom centered at 0.75 mm from the surface and connected to a microsyringe pump (70-2209, Harvard Apparatus, Holliston, MA). Microspheres were pumped through the phantom at flow rates of 25, 50, and 100 μL/min, which produced linear flow speeds of 15, 30 and 60 mm/s, respectively. A clear matching gel (#875465, McKesson Medical-Surgical, Inc., Richmond, VA) was applied between the phantom surface and the DiFC probe tip to minimize laser light specular reflection. We performed DiFC for the 6 microsphere concentration and speed combinations. This yielded DiFC detection rates from 1.5 to 19.5 per minute, approximately in the range of average count rates in MM and LLC mouse models. DiFC scanning was performed for 35 minutes in each case. Experiments were repeated in triplicate (N = 18).

### S.5 Simulated DiFC detections with Poisson processes.

As discussed in section 2.2, it is frequently assumed in the CTC literature that CTC detections should follow a Poisson process (3, 4). We simulated Poisson distributed DiFC detections in *silico* (so as to compare it to our measured in vivo data) using custom written Matlab (The Mathworks Inc., Natick, MA) code as follows. The time between CTC detections followed an exponential distribution given by:

$$P(t\ seconds\ between\ CTCs) = \lambda_{seconds} e^{-t\lambda_{seconds}}$$

Where $\lambda_{seconds}$ is the DiFC scan average number of cells per second. Detection times are generated until one exceeds the desired length of time of the scan. All detection times within that length of time then makeup the final simulated scan.

For Poisson Simulated DiFC data sets, we generated 35-minute data sets, where CTC detections were simulated with an exponential random number generator with mean rates of detection ($\lambda_{seconds}$) in the same range as the microsphere data sets.

For "Changing Mean" Simulated data sets, two 17.5 minute Poisson-distributed simulations were generated as described above, one with $\lambda_{seconds} = \lambda_1$ and the other with $\lambda_{seconds} = 2\lambda_1$. The $2\lambda_1$ detections are then appended to the end of the sequence of $\lambda_1$ detections to form a 35-minute scan. We generated 54 such data sets using with overall mean rates of detection in the same range as those measured in MM DXM mice. Of note, this process can also be considered a mixed Poisson process.

For "Merged Poissons" Simulated data sets, two 35-minute sequences of detections were generated, one with $\lambda_{seconds} = \lambda_1$ and the other with $\lambda_{seconds} = 2\lambda_1$. The sets of detection times were combined such that the Poisson-processes are concurrent (merged) over 35-minutes scans. We generated 54 such data sets.

### Supplementary References


1. R. Patil *et al.*, Fluorescence monitoring of rare circulating tumor cell and cluster dissemination in a multiple myeloma xenograft model in vivo. *J Biomed Opt* **24**, 1-11 (2019).
2. J. E. Fitzgerald *et al.*, Heterogeneity of circulating tumor cell dissemination and lung metastases in a subcutaneous Lewis lung carcinoma model. *Biomed Opt Express* **11**, 3633-3647 (2020).
3. A. L. Allan, M. Keeney, Circulating tumor cell analysis: technical and statistical considerations for application to the clinic. *J Oncol* **2010**, 426218 (2010).
4. A. G. Tibbe, M. C. Miller, L. W. Terstappen, Statistical considerations for enumeration of circulating tumor cells. *Cytometry A* **71**, 154-162 (2007).

**Supplementary Figures**

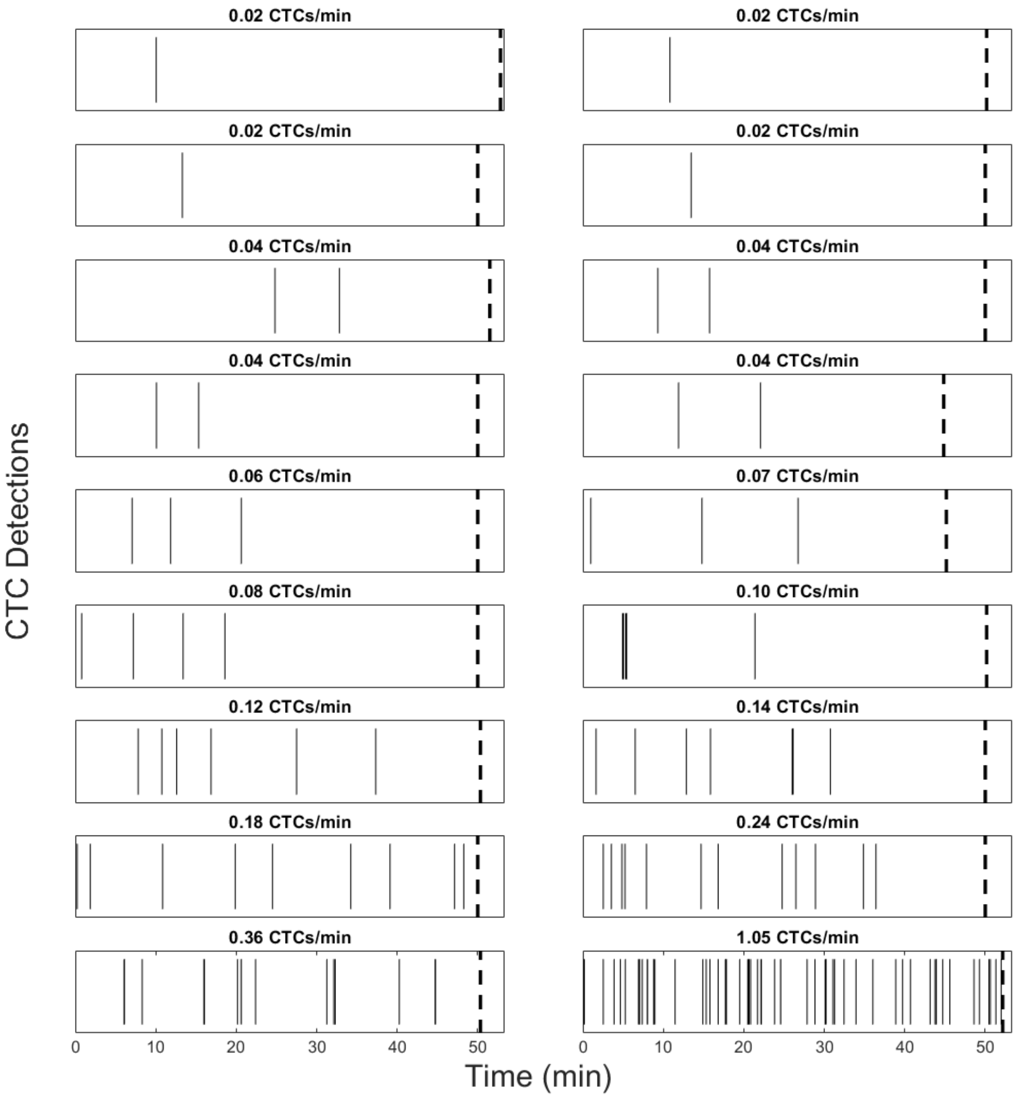


**Supplementary Figure S1.** Raster plots of 18 representative DiFC scans from LLC tumor bearing mice (from the "LLC data set"). Each solid vertical line represents a CTC detection. The dashed lines mark the end of each scan, which were of slightly different lengths. The plots are shown in ascending order of DiFC detection rate.

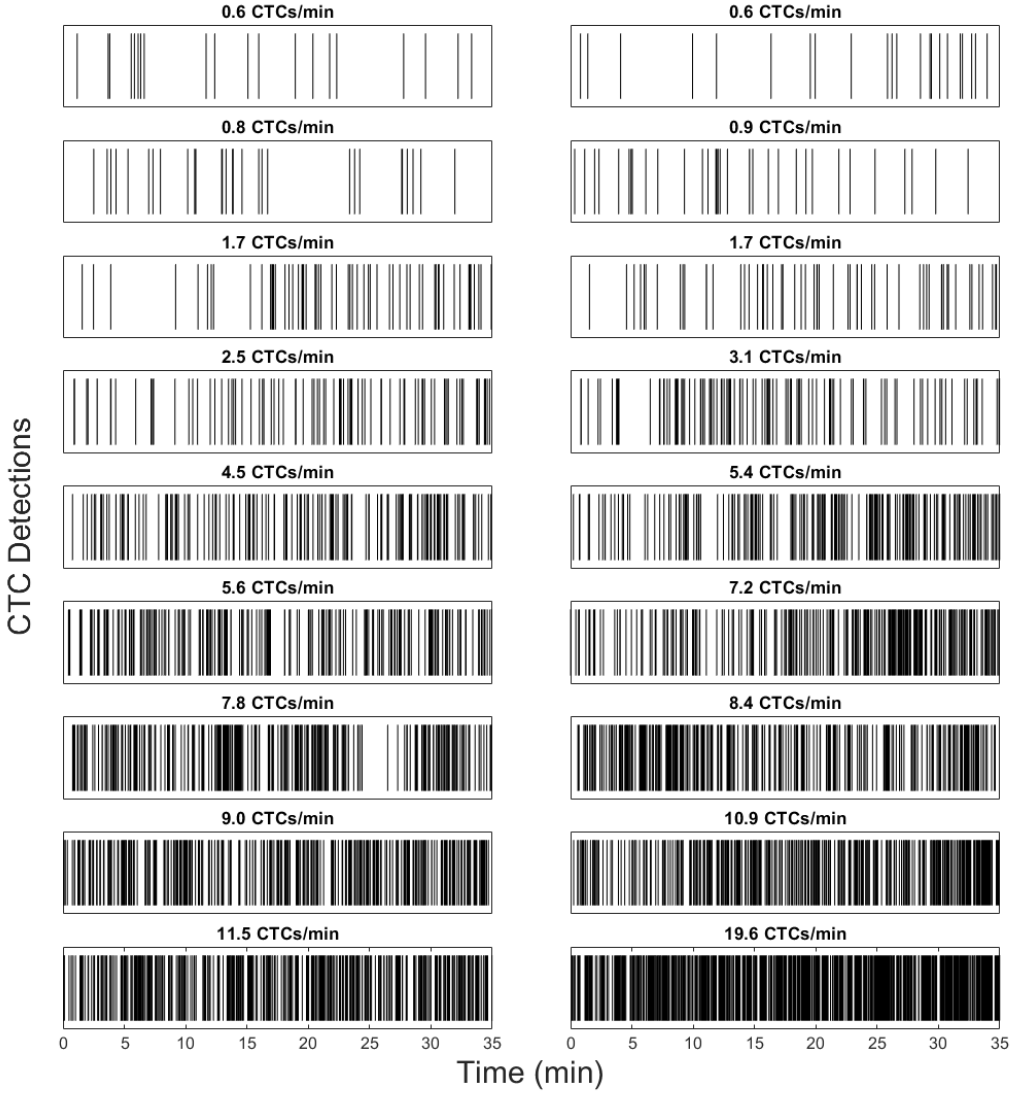


**Supplementary Figure S2.** Raster event plots for all 18 DiFC scans in the "MM 35-minute date set". Each vertical line represents a CTC detection. The plots are in ascending order of DiFC detection rate.

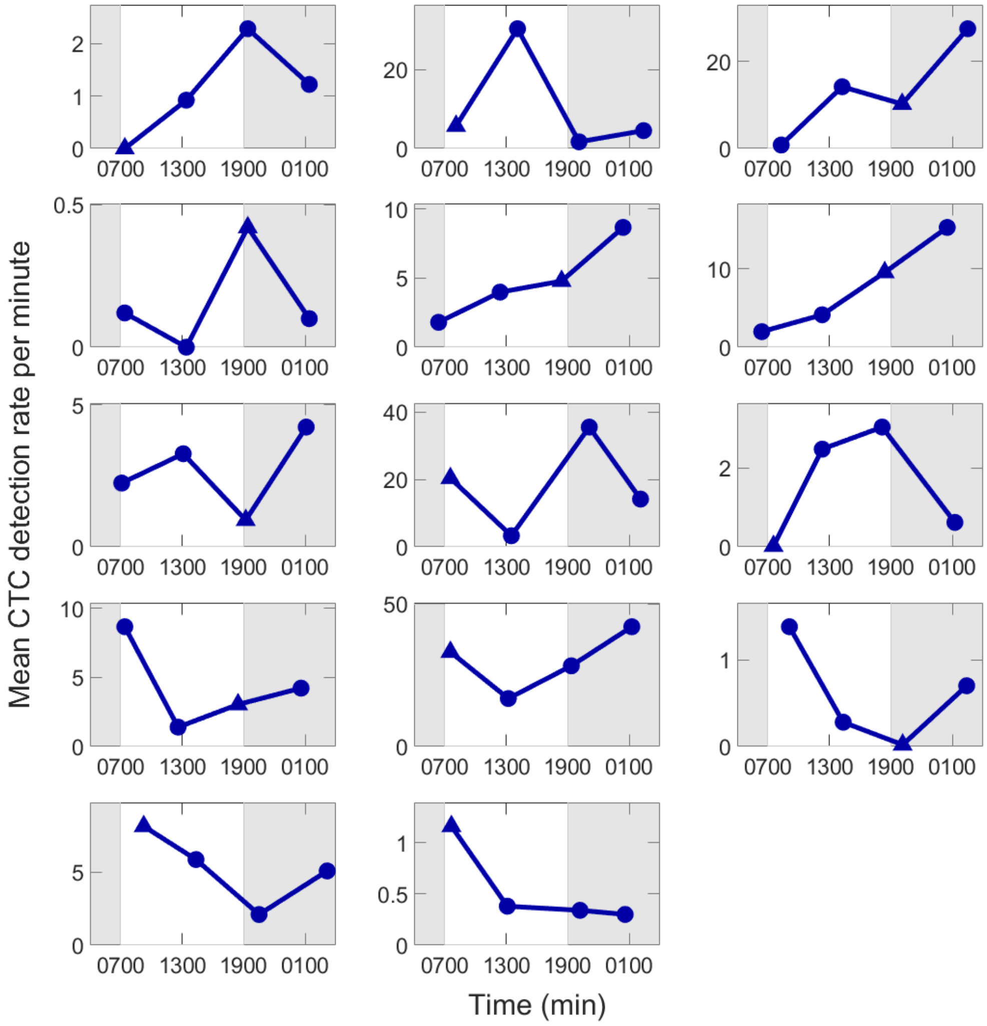


**Supplementary Figure S3.** Fluctuations in the mean CTC count rate over 24 hour periods for all DiFC scans the “MM 24-hour data set”. Triangle markers identify the first scan of the 24-hour cycle. N = 7 cycles started ~0700 (7 am) and N = 7 sessions began at ~1900 (7 pm). Animal housing followed a 0700 to 1900 light (white background) and 1900 and 0700 dark (grey background) light cycles.
